# Genome-wide characterization of the Fur regulatory network reveals a link between catechol degradation and bacillibactin metabolism in *Bacillus subtilis*

**DOI:** 10.1101/364588

**Authors:** Hualiang Pi, John D. Helmann

## Abstract

The ferric uptake regulator (Fur) is the global iron biosensor in many bacteria. Fur functions as an iron-activated transcriptional repressor for most of its regulated genes. There are a few examples where holo-Fur activates transcription, either directly or indirectly. Recent studies suggest that apo-Fur might also act as a positive regulator and, besides iron metabolism, the Fur regulon might encompass other biological processes such as DNA synthesis, energy metabolism, and biofilm formation. Here, we obtained a genomic view of the Fur regulatory network in *Bacillus subtilis* using ChIP-seq. Besides the known Fur target sites, 70 putative DNA binding sites were identified, and the vast majority had higher occupancy under iron sufficient conditions. Among the new sites detected, a Fur binding site in the promoter region of the *catDE* operon is of particular interest. This operon, encoding catechol 2,3-dioxygenase, is critical for catechol degradation and is under negative regulation by CatR and YodB. These three repressors function cooperatively to regulate the transcription of *catDE*, with Fur functioning as a sensor of iron-limitation and CatR as the major sensor of catechol stress. Genetic analysis suggests that CatDE is involved in metabolism of the catecholate siderophore bacillibactin, particularly when bacillibactin is constitutively produced and accumulates intracellularly, potentially generating endogenous toxic catechol derivatives. This study documents a role for catechol degradation in bacillibactin metabolism, and provides evidence that catechol 2,3-dioxygenase can detoxify endogenously produced catechol substrates in addition to its more widely studied role in biodegradation of environmental aromatic compounds and pollutants.

**Importance:** Many bacteria synthesize high affinity iron chelators (siderophores). Siderophore-mediated iron acquisition is an efficient and widely utilized strategy for bacteria to meet their cellur iron requirements. One prominent class of siderophores uses catecholate groups to chelate iron. *B. subtilis* bacillibactin, structurally similar to enterobactin (made by enteric bacteria), is a triscatecholate siderophore that is hydrolyzed to monomeric units after import to release iron. However, the ultimate fate of these catechol compounds and their potential toxicity have not been defined previously. Here, we performed genome-wide identification of Fur binding sites *in vivo* and uncovered a connection between catechol degradation and bacillibactin metabolism in *B. subtilis.* Beside its role in detoxification of environmental catechols, the catechol 2,3-dioxygenase encoded by *catDE* also protects cells from intoxication by endogeous bacillibactin-derived catechol metabolites under iron-limited conditions. These findings shed light on the degradation pathway and precursor recycling of the catecholate siderophores.

## Introduction

Iron is an essential micronutrient for most bacteria. It is required for many biological processes but can be toxic when present in excess. Various iron-mediated stress systems respond to changes in environmental iron availability (1, 2). Iron limitation induces acquisition systems to scavenge iron from the surroundings and activates systems to mobilize and prioritize iron utilization (3). Conversely, iron excess induces storage and efflux systems to maintain non-toxic levels of intracellular free labile iron (4, 5). These responses must be carefully coordinated by iron responsive regulators to ensure effective iron balance within the cell. The ferric uptake regulator (Fur) is the key regulator of iron homeostasis in many bacteria (6, 7). Fur monitors intracellular iron levels and regulates transcription of systems for iron uptake, utilization, storage, and efflux (7-9).

The Fur regulon has been characterized in many bacteria. In *B. subtilis*, the Fur regulon consists of an estimated 29 operons, many of which are involved in iron acquisition. These encode the biosynthesis machinery for the endogenous siderophore (bacillibactin, BB), and uptake systems for elemental iron, ferric citrate, BB, and various xenosiderophores that are secreted by other microbes (9). In general, Fur functions as an iron-activated transcriptional repressor for most of its regulon. Under iron replete conditions, Fur binds to its cofactor Fe^2+^ and the resulting holo-Fur binds to its target sites and represses transcription of its target genes; when iron is limited Fur loses its cofactor and apo-Fur dissociates from DNA leading to derepression of its regulon. Recent results reveal that the Fur regulon is derepressed in three sequential waves (3). As cells transition from iron sufficiency to deficiency, *Bacillus* cells (i) increase their capacity for import of common forms of chelated iron that are already in their environment such as elemental iron and ferric citrate, (ii) invest their energy to synthesize their own siderophore BB and produce high affinity siderophore-mediated import systems to scavenge iron, and (iii) activate a small RNA FsrA and its partner proteins to prioritize iron utilization (3).

In addition to its regulatory role as a transcriptional repressor, holo-Fur can also activate gene expression, either directly or indirectly (5, 10, 11). For instance, in *Escherichia coli* Fur positively regulates expression of the iron storage gene *ftnA* by competing against the H-NS repressor when iron levels are elevated (5), and *Listeria mononcytogenes* Fur activates the ferrous iron efflux transporter FrvA to protect cells from iron intoxication (10). Recent studies
suggested that apo-Fur may also act as a positive regulator (12) and, besides iron metabolism, the Fur regulon may expand into other biological processes such as DNA synthesis, energy metabolism, and biofilm formation (12-15). These findings motivated us to obtain a genomic view of the Fur regulatory network in response to iron availability in *B. subtilis.* Besides the known Fur target sites, 70 putative DNA binding sites were identified using chromatin immunoprecipitation coupled with high-throughput sequencing (ChIP-seq). Our attention was drawn to the binding site located in the promoter region of the *catDE* operon. This operon encodes a mononuclear iron enzyme, catechol 2.3-dioxygenase, which is critical for catechol degradation (16).

*B. subtilis* is a soil microbe. In its natural habitat, *B. subtilis* is exposed to many toxic aromatic compounds that are released from decaying plants, fungi, animals, and industrial wastes (17-19). Aromatic compounds are one of the most prevalent pollutants in the environment and the majority of them are oxidized to catechols before the benzene rings are cleaved (20). Catechols are persistent in the environment and can undergo a wide range of chemical reactions outside or within the cell: (i) complex formation with heavy metals, (ii) redox cycling, and (iii) production of reactive oxygen species (ROS) by reaction with metal ions and oxygen. These processes can damage DNA and protein and disrupt membrane potential (21).

Catechol dioxygenases have been widely studied for their initiating role in biodegradation of catechols, which are themselves common intermediates in the degradation of a wide variety of aromatic compounds (19). Our observation of a Fur-binding site preceding the *catDE* operon led us to hypothesize that, in addition to the external catechol stress, *B. subtilis* may encounter threats from the endogenous catechol compounds. In response to iron limitation, *B. subtilis* synthesizes the catechol-based siderophore BB and expresses siderophore-mediated uptake systems to overcome iron deficiency. BB, structurally similar to enterobactin primary found in Gram-negative bacteria (22), is synthesized through a non-ribosomal peptide synthetase (NRPS) assembly system (DhbACEBF) (23). BB is secreted by a major facilitator superfamily transporter YmfD (24), which is under regulation of the transcriptional activator Mta (24), a MerR family regulator of multidrug-efflux transporter system. BB chelates ferric iron with extremely high affinity (a pFe of ^~^10^−33^ M under biological conditions; (7)) and the resultant ferric-BB complex is then imported by the FeuABC-YusV system (25). Ferric-BB is subsequently hydrolyzed by the BesA esterase to release iron, which yield three BB-monomers (2,3-dihydroxybenzoate-glycine-threonine) (25). It is unknown whether the BB monomer is further processed, but this and derived catechol-containing compounds are potentially toxic and could affect cell fitness.

In this study, we demonstrate that holo-Fur functions as a repressor and works cooperatively with two other regulators, CatR and YodB, in regulating transcription of the *catDE* operon. This operon is induced upon catechol stress or iron limitation, and strongly induced when both conditions are present. Furthermore, accumulation of endogenous BB-derived catechol compounds triggers cell lysis and CatDE is required to alleviate the toxicity. These findings suggest that CatDE is involved in metabolism of the triscatecholate siderophore BB and reveal a link between catechol degradation and bacillibactin metabolism in *B. subtilis*.

## Results and discussion

### Genome-wide identification of Fur-binding sites by ChIP-seq

A recent study suggested that under anaerobic conditions *B. subtilis* Fur might regulate genes beyond its previously defined regulon (15). Moreover, our previous genome-wide studies of Fur regulation were focused on those genes (as monitored by microarray analysis) that were derepressed in both a *fur* mutant and in response to iron depletion. Since Fur might also act to activate gene expression, and some targets might not have been represented in the microarray (which was focused on annotated ORFs), we here chose to take an unbiased view towards defining those sites bound to Fur *in vivo* under both iron replete and iron deficient conditions using ChIP-seq. To modulate intracellular iron levels, we employed a high-affinity Fe^2+^ exporter FrvA from *L. monocytogenes* to impose iron starvation, as described previously (3, 10). *Bacillus* wild-type cells (with C-terminal FLAG-tagged Fur at its native locus and an IPTG-inducible ectopic copy of *frvA* integrated at *amyE* locus) were harvested at 0 and 30 min after IPTG induction to study Fur-dependent regulation under iron sufficient and deficient conditions, respectively (see details in “Materials and Methods”).

### Fur-dependent binding under iron deficient conditions

ChIP-seq analysis identified 89 and 27 reproducible Fur-binding sites (signal to noise ratio, S/N ≥1.5) under iron sufficient and deficient conditions, respectively (Fig. 1 and Table S3-S4). Most of the binding sites (22 out of 27) occupied under iron deficient conditions overlap with those under iron sufficient conditions, so the total number of binding sites are 94 (Fig. 1B). These sites may represent sites bound by holo-Fur that are of particularly high affinity and slow dissociation rate, or possibly sites that can be occupied *in vivo* by apo-Fur.

**Fig. 1.**
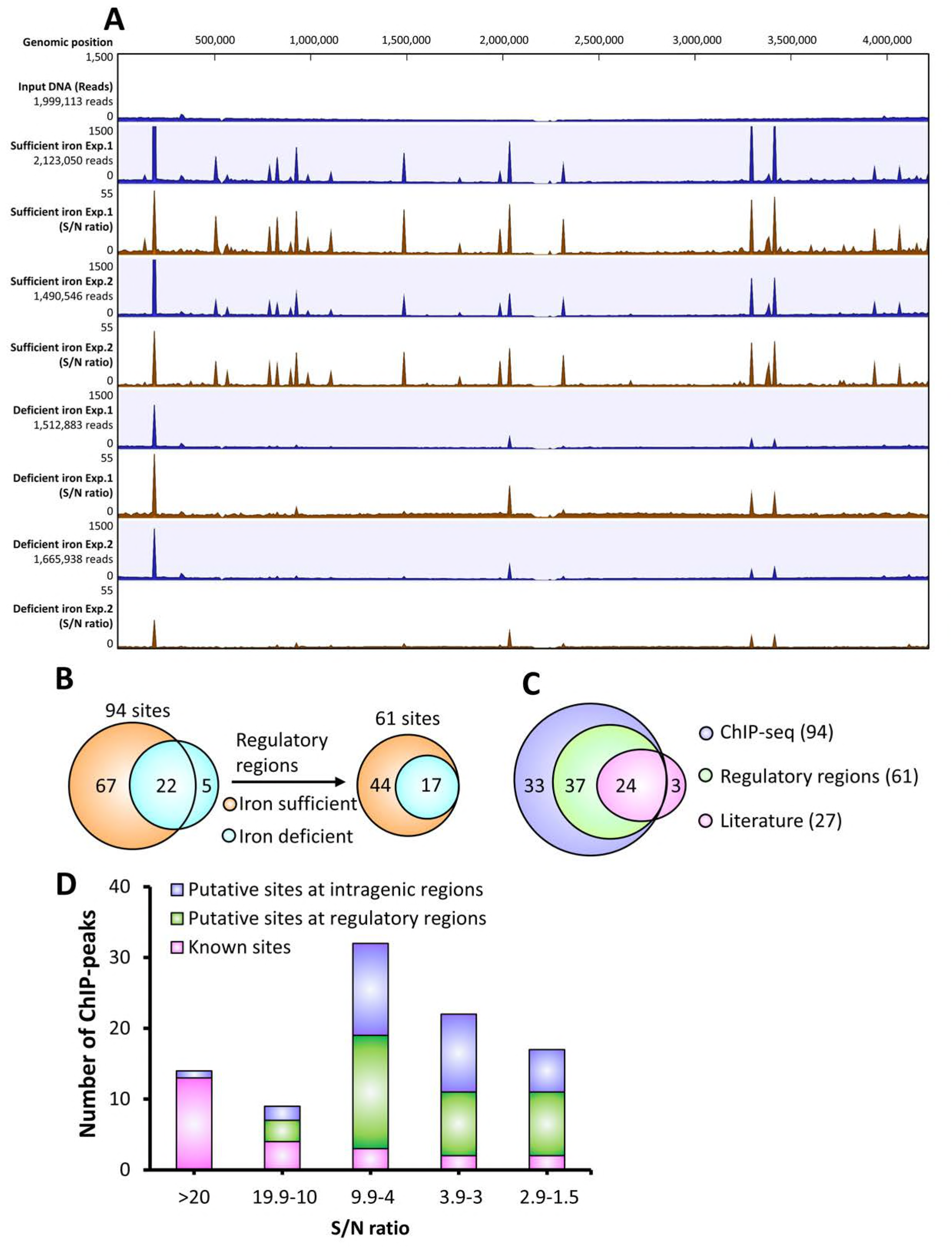
An overview of Fur-binding profiles across the *B. subtilis* genome under varied iron conditions. A. ChIP-seq data of *B. subtilis* Fur-dependent binding under iron sufficient and deficient conditions. Two biological replicates were included for each growth condition (Exp. 1 and Exp. 2). Reads are in blue with maximum set at 1500 for all the ChIP peaks. The total reads for each sample are included on the left side. S/N denotes signal to noise ratio for peak calling as shown in brown.
B. Most of ChIP peaks (22 out of 27) identified under iron deficient conditions overlap with those detected under iron sufficient conditions. The majority of peaks (61 out of 94) are located in regulatory regions.
C. Most of the known Fur binding sites from the literature (24 out of 27) are identified by ChIP-seq with three exceptions. Among all the new identified Fur binding sites, 33 sites are located in intragenic regions while 37 sites are located in regulatory regions.
D. Distribution of ChIP peaks relevant to S/N ratio. The highest S/N ratio of each ChIP peak among all four samples was used for this analysis.

Five ChIP peaks are specific to Fur under iron deficient conditions, and could represent authentic apo-Fur specific sites (Fig. S1 and Table S4). One of these sites is located in the promoter of the *S477*-*ykoP* operon (Table S3), which encodes a possible regulatory RNA S477 and a protein YkoP with unknown function. This operon has been implicated to be under negative regulation of multiple regulators including Fur (9), ResD (15), NsrR(15), and Kre (26). However, a statistically significant ChIP-peak (P-value ≥ 0.05) was only detected in one of the biological replicates (Table S3), indicating that the regulatory role of Fur at this site is uncertain. Four other apo-Fur specific sites are located in intragenic regions and Fur occupancy at these sites is fairly low (Fig. S1 and Table S4). The physiological significance of Fur-binding at these sites is unclear. However, the ChIP peak located inside *ylpC* (also known as *fapR*) is very close to 5’-end of the gene (Fig. S1D). FapR functions as a global regulator of fatty acid biosynthesis and is well conserved in Gram-positive bacteria (27). The *fapR* gene is under dual regulation: negative regulation by FapR itself (28) and positive regulation by the quorum sensing regulator ComA (29). Interestingly, transcriptome data show that expression of *fapR* is ^~^4-fold upregulated in a *fur* null mutant compared to WT (9), suggesting that *fapR* might be under negative regulation of Fur under iron deficient conditions. Overall, our results suggest that apo-Fur has a limited role, if any, in gene regulation in *B. subtilis.* However, the connection between fatty acid biosynthesis and iron homeostasis deserves further investigation.

### Known Fur target sites identified by ChIP-seq

Most of the previously defined Fur target sites (24 out of 27) were detected *in vivo* by ChIP-seq analysis (Fig. 1 and Table S3), which validated the modified method of ChIP-seq in *B. subtilis* (see details in “Materials and Methods”) and further confirmed the Fur binding sites characterized by *in vitro* DNase 1 footprinting and transcriptome analysis (9). However, Fur occupancy at the promoter sites of *pfeT*, *yfkM*, and *ydhU/2*-*ydhU/1* was undetectable. The gene *pfeT* encodes a Fe^2+^-efflux transporter and its expression is activated by Fur only under excess iron conditions (30), which is likely why we were unable to detect Fur binding at this site under the conditions tested. The gene *yfkM* encodes a general stress protein that is under regulation of SigB, and *ydhU2* and *ydhU/1* are two inactive pseudogenes. The prior study only detected very weak Fur binding at the promoter sites of *yfkM* and *ydhU/2*-*ydhU/1 in vitro* and the Fur box identified at these two sites are only 10 out of 15 bases matching to the minimal 7-1-7 consensus sequence (9, 31). These results, together with the lack of measurable Fur occupancy *in vivo*, suggest that Fur may not play a significant role in regulation of these genes.

### ChIP peaks located in intragenic region

Many of the putative Fur binding sites identified by ChIP-seq (29 out of 70 sites) are located in intragenic regions (Table S4). Expression of most of these genes are not regulated by Fur (comparing mRNA levels between a *fur* mutant and WT) as monitored by microarray analysis (9) and qPCR (Table S4 and Table S5), indicating an apparent lack of physiological relevance for these sites. We then evaluated the possible involvement of some putative targets in iron homeostasis, either iron intoxication or limitation. Among the five targets tested (the ones associated with high Fur occupancy i.e. *ppsB*, *gidA*, *tufA*, *ybaC* and *yycE*), none of them showed sensitivity to high iron; only the *gidA* null mutant showed modest sensitivity to dipyridyl compared to WT. The gene *gidA* encodes a tRNA uridine 5-carboxymethylaminomethyl modification enzyme and its role in iron homeostasis in currently under further investigation.

Another notable candidate for a functional intragenic Fur-binding site resides within *ppsB*, the second gene in a long operon encoding a non-ribosomal peptide synthetase (NRPS) that synthesizes the antibacterial compound plipastatin (a lipopeptide closely related to fengycins) (32). This peak has the highest Fur occupancy among all the newly identified sites (Table S4). A promoter and two transcription start sites in opposite directions were assigned overlapping this ChIP peak (33), suggesting that transcription initiates internally to *ppsB*. In addition, a putative Fur box was identified within this peak area (11 out of 15 bases matching to the consensus sequence) (Table S5). The transcriptome data showed that expression of *ppsB* is ^~^3-fold downregulated in a *fur* null mutant compared to WT, although this result was not confirmed by qPCR (Table S5). The *ppsB* null mutant showed no significant sensitivity to either iron intoxication or limitation compared to WT (Table S5). However, numerous studies document an important stimulatory role for iron in lipopeptide production in *Bacilli* (34, 35), and lipopeptides can chelate metal ions (36) and likely iron. Together, these results lead us to speculate that transcripts encoded within *ppsB* may be regulated by Fur, and perhaps function to coordinate plipastin synthesis with iron status.

### ChIP peaks located in regulatory regions

Among the 70 putative Fur target sites identified by ChIP-seq analysis, 37 are located in regulatory regions (Fig. 1C-D, Table S4). Most of these sites bind Fur under iron sufficient conditions, although Fur also binds at some sites in at least one of the biological replicates under iron deficient conditions (Table S4). At least twelve of these have good Fur boxes matching to the minimal 7-1-7 consensus sequence (Table S5). Interestingly, Fur binds to the promoter region of *gntR* in an iron-independent manner: Fur occupancy at this site remained about the same level under either iron deficient or sufficient conditions (Table S4).

To evaluate the involvement of these putative Fur targets in iron homeostasis, we constructed deletion mutants of the top twelve candidates and carried out assays to test their sensitivity to high levels of iron and dipyridyl (Table S5). None of them showed significant sensitivity to high iron. Five of them showed moderate sensitivity to dipyridyl, including *cspB*, *yhcJ*, *catD*, *narJ*, and *yybN* (Table S5). Transcriptome data suggest that expression of *cspB*, *catD*, and *narJ* might be under regulation of Fur (3). The expression of *catD* is upregulated (3.3) whereas that of *cspB* (0.3) and *narJ* (0.2) is downregulated in a *fur* null mutant compared to the WT strain (Table S5). The regulatory role of Fur in *catD* and *narJ* was further confirmed by mRNA quantification using qPCR (Table S5). Here, the bicistronic operon *catDE* was subject to further investigation to understand its physiological role in iron homeostasis.

### Fur binds to the regulatory site of *catDE* operon under sufficient iron conditions

A reproducible ChIP peak was identified in the promoter region of *catDE* operon under sufficient iron conditions. The signal to noise ratio for peak calling is relatively high in both independent replicates (10.1 for Exp.1 and 14.1 for Exp. 2; Fig. 2 and Table S4), and the ChIP DNA enrichment at this site, compared to the input DNA control, is statistically significant (P-value is 3.5×10^−24^ for Exp. 1 and 1.2×10^−44^ for Exp.2; Table S4). The *catDE* operon encodes a putative catechol 2,3-dioxygenase that requires Fe^2+^ as its cofactor (16). CatE showed 2,3-dioxygenase activity *in vitro* and is essential for viability in the presence of catechol (16, 37).

**Fig. 2.**
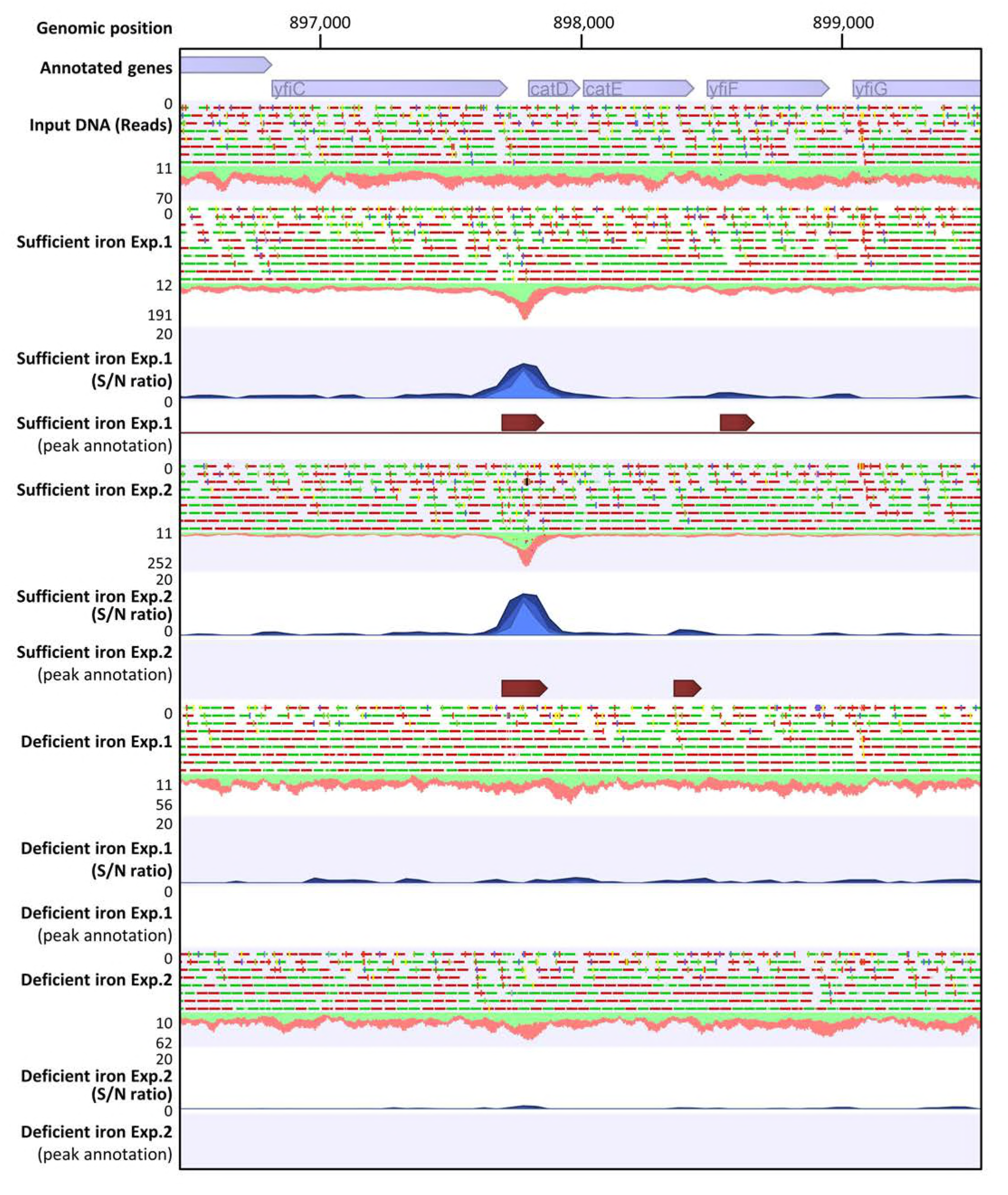
Fur binds to the promoter region of *catDE* operon under iron replete conditions. A zoom-in example of Fur binding at the *catDE* operate site identified by ChIP-seq. Two biological replicates were included as Exp.1 and Exp.2 for each condition. A single peak was annotated at this region for both replicates under iron sufficient conditions. The peak length was 328 bp for Exp. 1 and 355 bp for Exp. 2. No peaks were annotated at this region for either replicate under iron deficient conditions. S/N denotes signal to noise ratio for peak calling.

We tested the sensitivity of either single (*catD* or *catE*) or double mutant (*catDE*) strains to catechol toxicity using both disk diffusion and bioscreen growth assays. Our results confirmed that they are both involved in catechol detoxification (Fig. S2). Both assays were performed in Belitsky minimal medium because we noticed that the catechol toxicity is significantly diminished in LB medium (Fig. S3), probably due, at least in part, to metal-catechol complexes formed with Fe, Cu, Mn, and other divalent metal ions that are present in LB medium (Fig. S4).

### Regulation of the *catDE* operon by three regulators

The *catDE* operon is under negative regulation of CatR, a MarR/DUF24 family transcription regulator that senses catechols, and YodB, a regulator of genes important for quinone and diamide detoxification (16). In addition, a Fur box (13 out of 15 bases match to the minimal 7-1-7 consensus sequence; (31)) is located downstream of the transcription start site (Fig 3A), suggesting that Fur may act as a repressor. Indeed, qPCR measurements indicate that expression of *catD* was upregulated ^~^4-fold in the *fur* null mutant compared to wild-type cells (Fig. 3C), consistent with the previous transcriptome analysis (9).

**Fig. 3.**
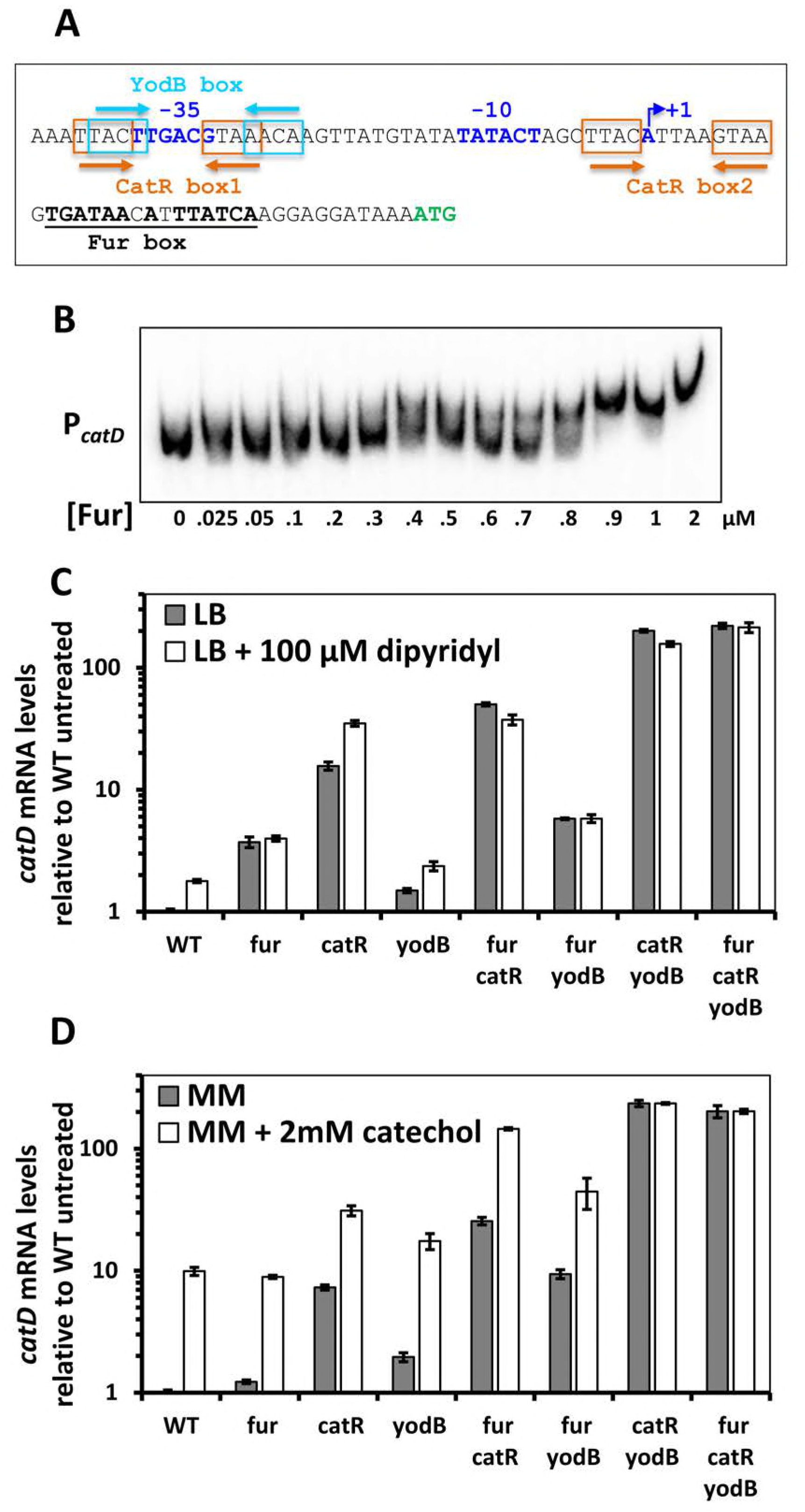
Regulation of the *catDE* operon by three transcription factors. A. The promoter sequence of the *catDE* operon: –10, –35, and the transcriptional start site (+1) are highlighted in blue; the two CatR boxes are indicated by orange arrows; the YodB box is indicated by light blue arrow; the Fur box is underlined.
B. Electrophoretic mobility shift assay (EMSA) was carried out to determine Fur-DNA binding affinity to the promoter region of the *catDE* operon. Experiments were repeated three times and a representative image is shown.
C. The *catD* mRNA levels were compared among different strains grown in LB medium without or with 100 μM dipyridyl using qPCR.
D. The *catD* mRNA levels were compared among different strains grown in Belitsky minimal medium without or with 2 mM catechol using qPCR. The 23S rRNA gene was used as an internal control for both C and D.

We used electrophoretic mobility shift assays (EMSA) to determine the affinity of Fur for the *catDE* operator site *in vitro.* Surprisingly, unlike the very high affinity (K_d_ ^~^0.5-5.6 nM) observed for Fur binding to most of its known target sites (3), the affinity of Fur for the *catDE* promoter is quite low (K_d_ ^~^0.7 μM). However, this affinity is comparable to that of either CatR or YodB binding to the same promoter region (16), and all appear to bind rather weakly when tested individually. This suggests that perhaps these three regulators interact with one another and bind to the promoter site cooperatively.

To dissect the cooperativity among the three regulators *in vivo*, a genetic study was performed using single, double, and triple deletion mutants of these three regulators and *catD* mRNA levels were quantified under various stress conditions: iron sufficiency, iron limitation, and catechol stress. Consistent with the prior study (16), CatR functions as the major regulator and YodB plays a minor role in regulation of the *catDE* operon (Fig. 3C-D). When only one regulator is present (in the double mutants), YodB repression (in *fur catR* double mutant) resulted in a ^~^4-fold reduction in mRNA compared to the full derepression observed in the triple mutant (*fur catR yodB*). CatR is the major repressor, and accounts for a ^~^38-fold reduction of *catD* expression (comparing the *fur yodB* double mutant to the *fur catR yodB* triple mutant). Interestingly, the *catD* mRNA level in *catR yodB* double mutant is comparable to that in the triple mutant, indicating that Fur plays a negligible regulatory role when both CatR and YodB are absent, and Fur binding at this site *in vivo* may require, or facilitated by, the other two regulators.

### Either CatR or YodB facilitates Fur binding at the promoter site of *catDE*

To further explore whether CatR and/or YodB facilitate Fur binding *in vivo*, Fur occupancy at the promote site of *catDE* was evaluated using ChIP-qPCR. No noticeable change of Fur occupancy was observed in the *catR* single mutant compared to wild-type cells, while Fur occupancy increased significantly in *yodB* single mutant (Fig. 4A), indicating that Fur interacts with CatR more efficiently when YodB is absent. This is consistent with the expression data, which showed that *catD* was induced by dipyridyl in the *catR* single mutant to a level comparable to that observed in the *fur catR* double mutant, whereas it was only partially derepressed by dipyridyl in the *yodB* single mutant compared to the *fur yodB* double mutant (Fig. 3C). Interestingly, Fur occupancy decreased significantly when both regulators are absent in the *catR yodB* double mutant (Fig. 4A). As expected, neither CatR or YodB affected Fur occupancy at the promoter of *dhbA* (Fig. 4B). These results suggest that CatR, and to a lesser extent YodB, facilitate Fur binding at the promoter region of *catDE.* Similarly, NsrR and ResD have been reported to facilitate Fur binding at a minority of co-regulated sites under anaerobic conditions; at most common sites binding was competitive (15). At the apparently cooperative sites (*ykuN*, *fbpC* and *exlX/yoaJ*), Fur appeared to facilitate binding of NsrR and/or ResD.

**Fig. 4.**
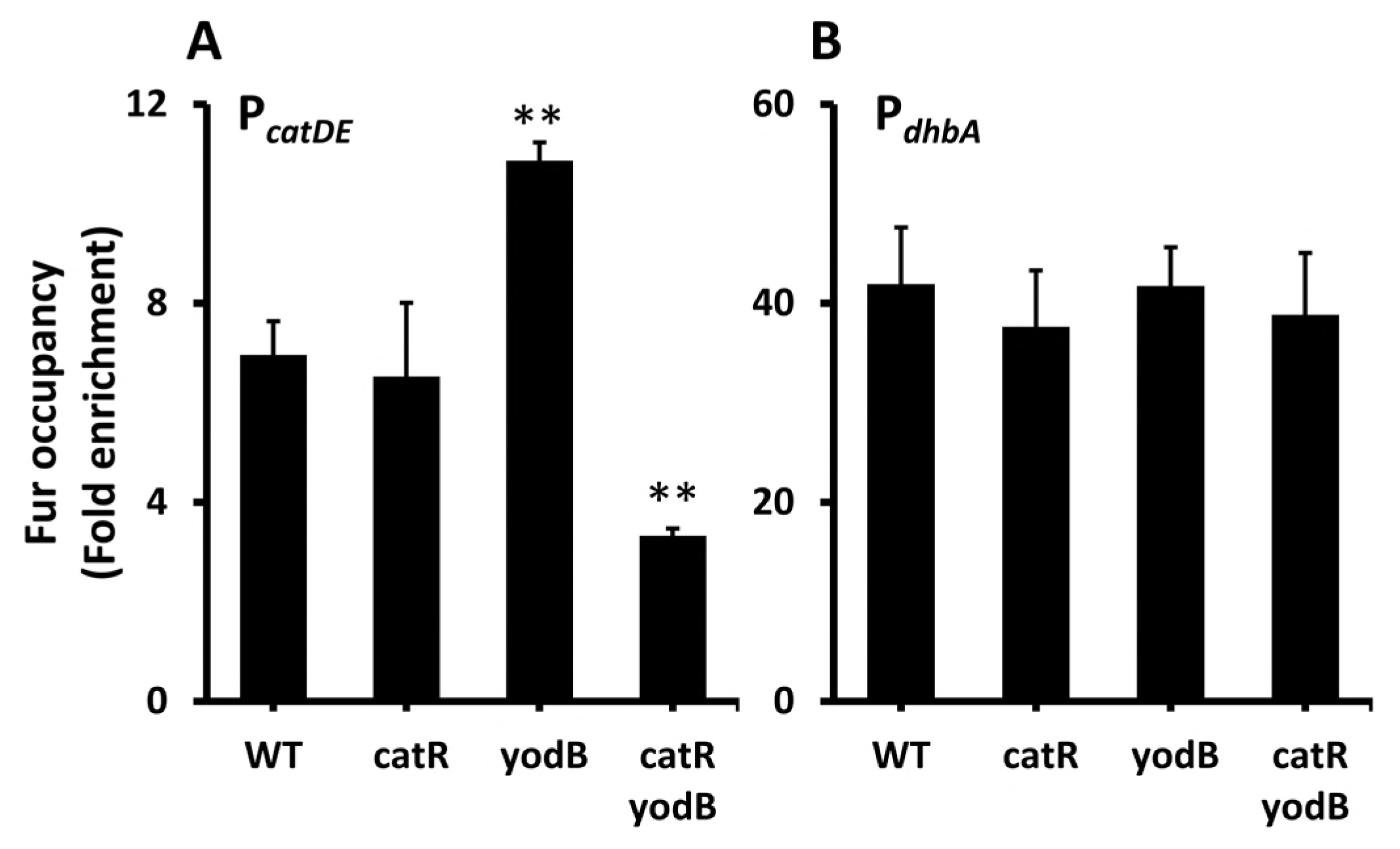
Either CatR or YodB facilitates Fur binding at the promoter site of *catDE.* Fur occupancy was evaluated by chromatin immunoprecipitation (ChIP) using anti-FLAG antibodies. Coimmunoprecipitated DNA was quantified by qPCR. DNA enrichment was calculated based on the input DNA (1% of total DNA used for each ChIP experiment). The Data are presented as the fold enrichment of Fur occupancy at the promoter sites of *catDE* (A) and *dhbA* (B) (mean ± SD; n = 3). Significant differences between wild type and mutant strains are indicated: ^∗∗^P < 0.01. No significant DNA enrichment was observed for *gyrA* that serves as a negative control.

### CatDE is involved in bacillibactin metabolism

The *B. subtilis* wild type strain (168) and its derivatives does not normally synthesize BB due to a null mutation in the *sfp* gene (*sfp^0^*) encoding the phosphopantetheinyl transferase required for activation of the Dhb NRPS complex. Strains lacking functional Sfp secrete a mixture of 2,3-dihydroxybenzoate (DHBA) and its glycine conjugate (DHBG), collectively known as DHB(G). We use both an *sfp^0^* strain and an isogenic strain with a corrected *sfp* gene (*sfp*^+^) that produces BB for our iron homeostasis studies.

The expression of *catDE* is induced ^~^3-fold in both *sfp^0^* and *sfp*^+^ strains upon iron depletion imposed by dipyridyl (Fig. 3 and Fig. 6A-B), suggesting that CatD and/or CatE may be involved in iron homeostasis. To test this idea, dipyridyl sensitivity of single (*catD* and *catE*) and double (*catDE*) mutant strains in both *sfp^0^* and *sfp*^+^ backgrounds was evaluated using a disk diffusion assay. Indeed, both CatD and CatE play important roles in times of iron limitation since mutants were more strongly growth inhibited in the presence of the iron chelator dipyridyl (Fig. 5C-D). We therefore hypothesized that the enzymatic activity of CatDE may be required to detoxify BB-derived catechol compounds produced upon iron limitation. When iron is limited, BB is secreted into the environment to acquire iron and the Fe^3+^-BB complex is imported back into the cell through the FeuABC-YusV system. Iron is released and BB is cleaved into BB monomers (2,3-dihydroxybenzoate-gly-thr), and perhaps further processed into catechol derivatives, which may require CatDE for detoxification.

**Fig. 5.**
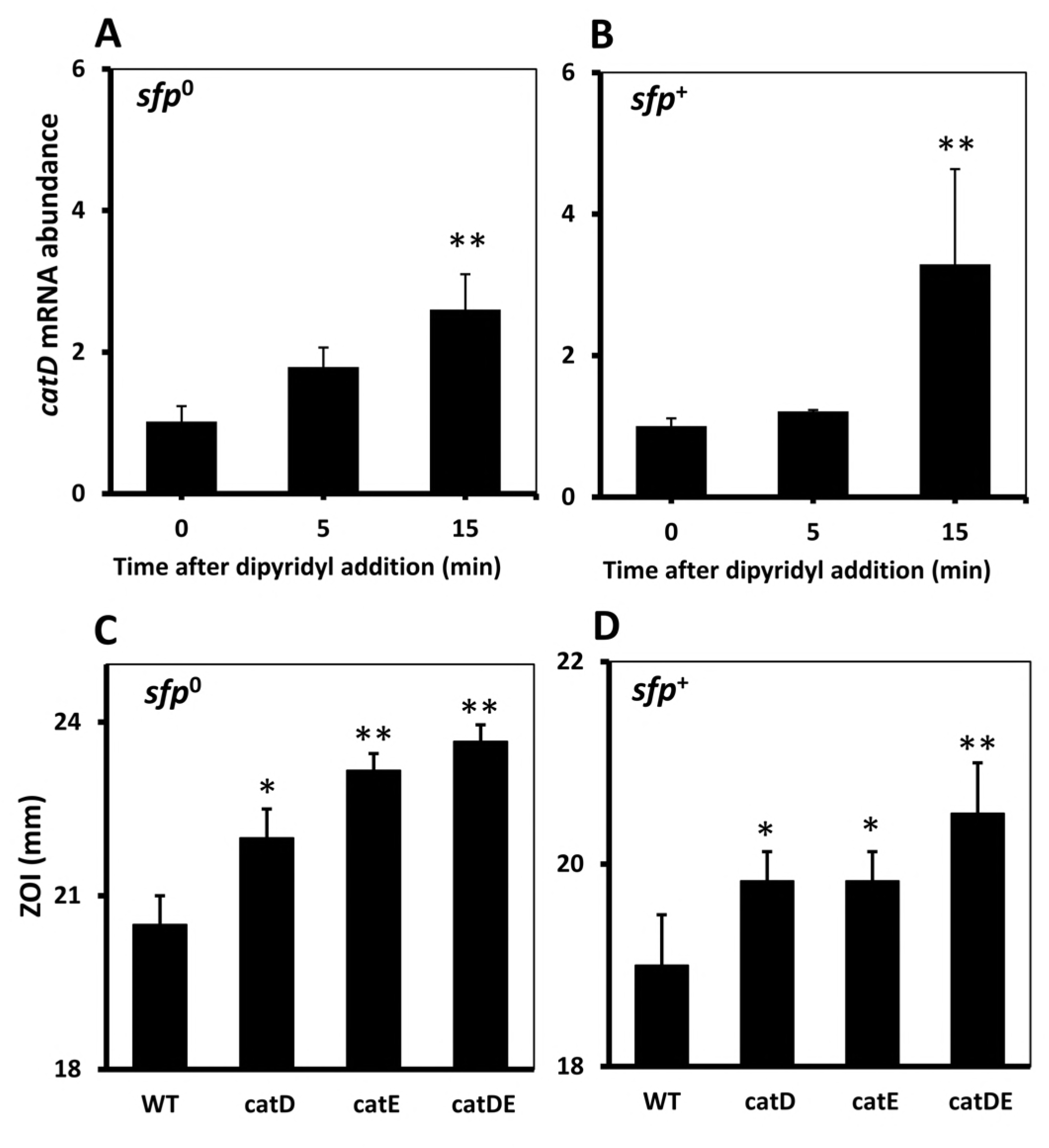
*catDE* is induced upon iron depletion and plays an important role in iron homeostasis. (A,B) Expression of *catD* was monitored in wild-type cells (A, *sfp^0^;* B, *sfp*^+^) grown in LB medium before (0 min) and after treatment of 100 μM dipyridyl. Significant difference between treated and untreated groups is indicated as ^∗∗^P < 0.01. (C,D) Dipyridyl sensitivity of WT, single (*catD* and *catE*), and double (*catDE*) mutant strains in both *sfp^0^* (C) and *sfp*^+^ (D) backgrounds was evaluated using a disk diffusion assay. The data are expressed as the diameter (mean ± SEM; n = 3) of the inhibition zone (mm). Significant differences between wild type and mutant strains are indicated: ^∗^P<0.05 and ^∗∗^P < 0.01.

We employed a genetic approach to evaluate the involvement of CatDE in BB metabolism. We used a *fur ymfD* double mutant in which BB is constitutively produced and accumulates intracellularly due to the loss of the YmfD BB exporter. We then asked whether the *catDE* operon is important for growth under these conditions. No growth defects were noticeable in the *fur ymfD catDE* quadruple mutant compared to the *fur ymfD* double mutant in the first 6h, however, dramatic cell lysis was observed in the quadruple mutant afterwards, while the double mutant continued growing (Fig. 6A). We infer that the *catDE* operon is critical for maintaining cell fitness when BB-derived catechol compounds accumulate intracellularly. Indeed, introduction of a *dhbA* null mutation to the quadruple mutant significantly rescued the cell lysis phenotype (Fig. 6A).

**Fig. 6.**
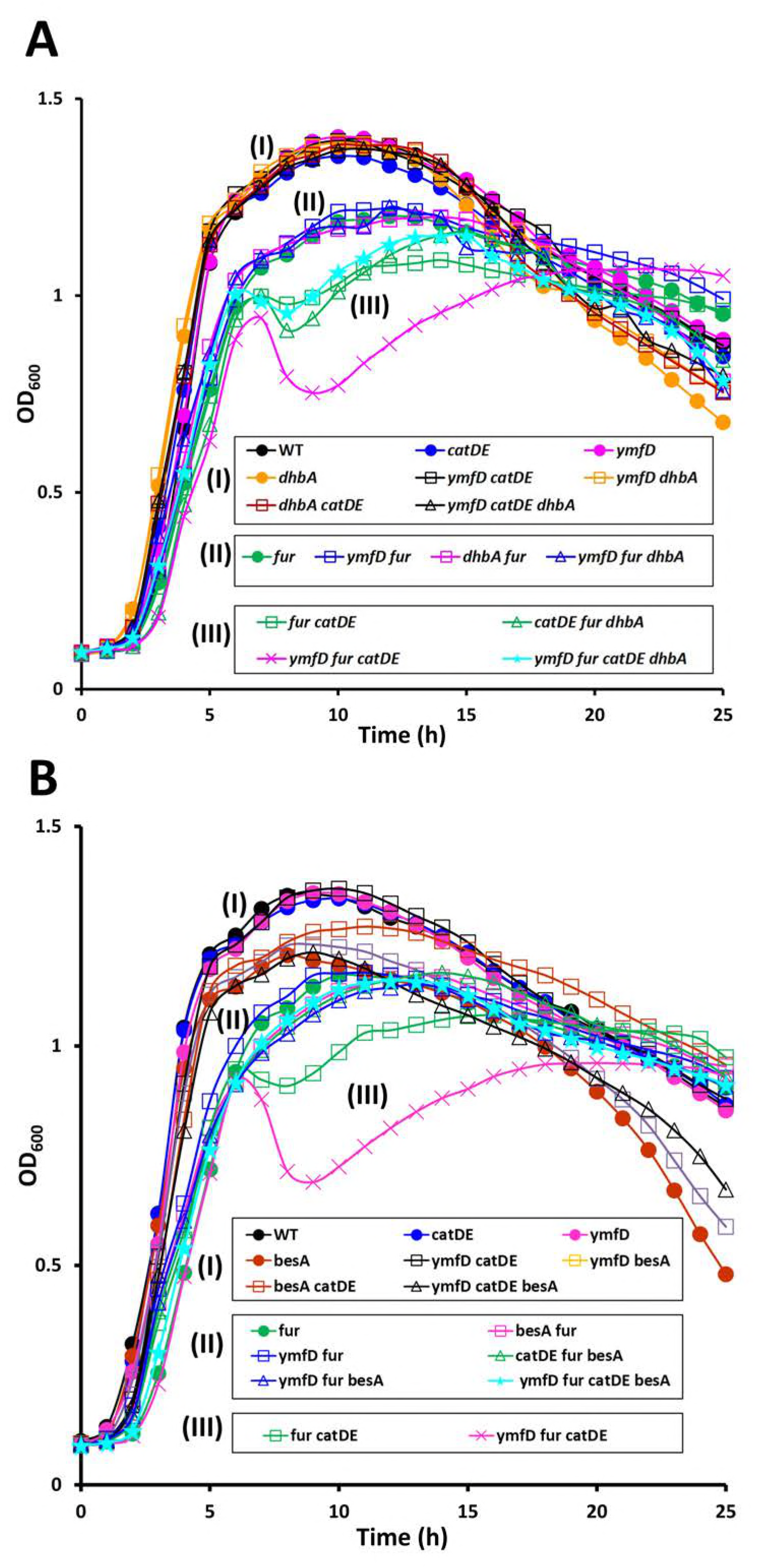
CatDE is involved in BB metabolism. (A, B) Representative growth curves are shown for strains that constitutively produce BB (*fur* mutation) and accumulate BB intracellularly (due to loss of the BB exporter YmfD). To check whether CatDE is required for BB metabolism growth of wild type (*sfp*^+^) and its derived mutant strains was monitored in LB medium for 25 h. Experiments were performed three times with three biological replicates each time.

**Fig. 7.**
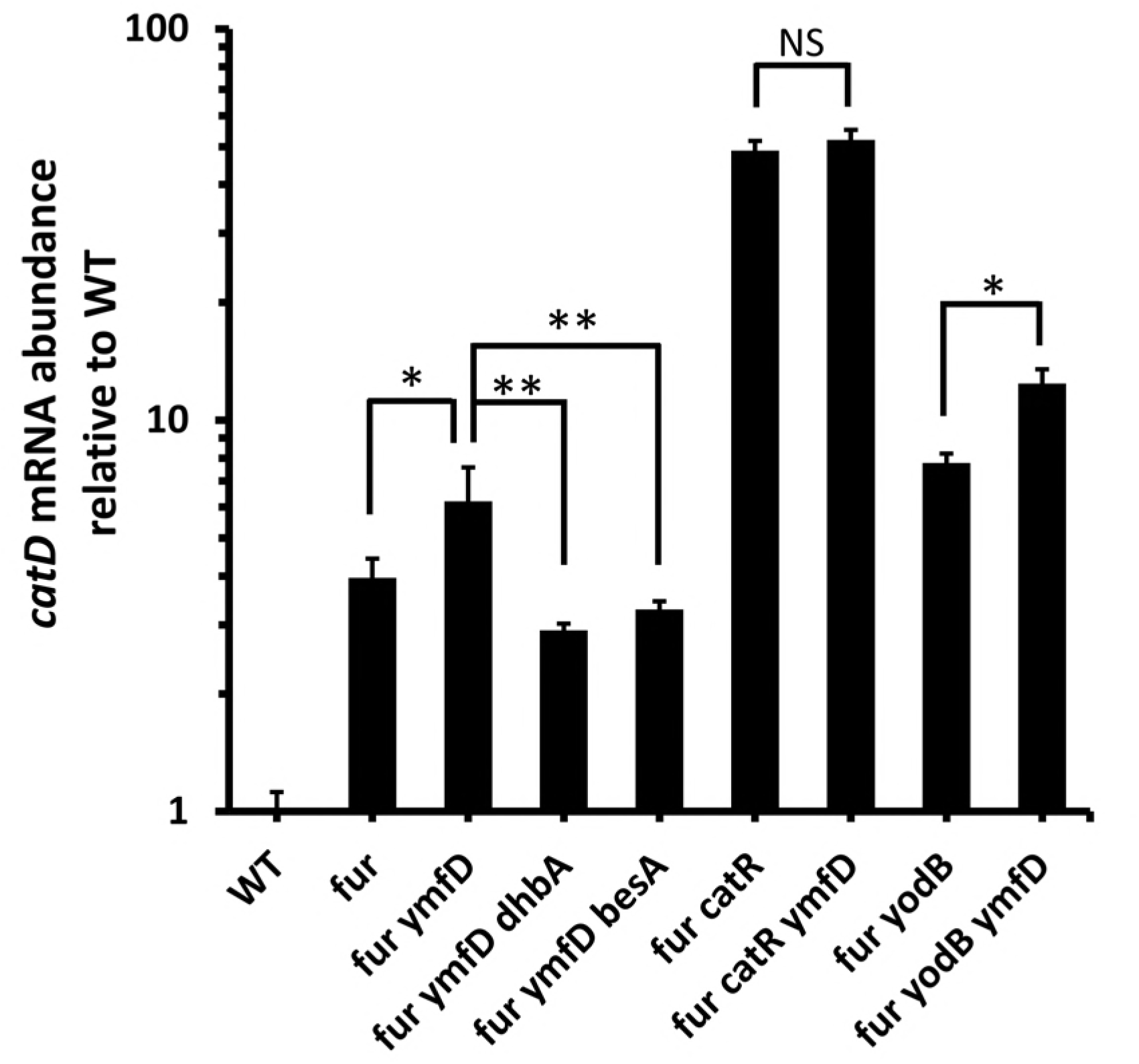
Accumulation of intracellular BB-derived catechol induces *catD* expression. The mRNA expression levels of *catD* was evaluated in WT (*sfp*^+^) and its derived mutants grown in LB medium to an OD_600_ of ^~^0.4. The 23S rRNA was used as an internal control. Significant differences are indicated as ^∗^P < 0.05 and ^∗∗^P<0.01. NS denotes not significant.

Once BB is imported back into the cytosol, it is hydrolyzed by the BesA esterase to release the chelated iron and the siderophore is cleaved into three BB monomers. To understand whether the cell lysis defect is due to accumulation of BB or BB monomer, we introduced a *besA* null mutation to the quadruple mutant (*fur ymfD catDE*). In the absence of BesA, cell lysis defect was no longer observed and cells grew almost as well as the *fur ymfD* double mutant (Fig. 6B). It is unknown whether or how BB monomers are further processed. Nevertheless, the BB monomer and perhaps derivative catechol compounds clearly require CatDE for detoxification.

### Accumulation of intracellular BB-derived catechol induces *catD* expression

Since intracellular BB-derived catechol compounds can compromise cell fitness, we wished to determine if they might also serve as inducers of the *catDE* operon. To test this, we compared *catD* mRNA levels in the *fur ymfD* double mutant and the *fur* single mutant. Indeed, *catD* expression is increased in the double mutant (defective for BB efflux) compared to the *fur* single mutant. This induction is specifically due to accumulation of intracellular BB-derived catechols since deletion of either *dhbA* or *besA* abolished induction. To understand which regulator is responsible for this induction, we monitored the *catD* mRNA levels in *fur catR* and *fur yodB* double mutants. When both Fur and CatR were absent, induction was no longer evident, suggesting that YodB does not respond to the accumulating catechols. In contrast, when both Fur and YodB were absent, a similar level of induction was observed. These results suggest that intracellular accumulation of BB-derived catechol metabolites can lead to at least partial inactivation of the CatR repressor, thereby leading to induction of the *catDE* operon. Since Fur and CatR bind cooperatively *in vivo*, this system is tuned to respond sensitively to the accumulation of catechol compounds (sensed by CatR) during times of iron starvation (sensed by Fur).

### Fur occupancy on the *catDE* operator site

The Fur target genes are derepressed in three sequential waves upon iron depletion (3), which provides insights into the distinct roles of the Fur targets in iron metabolism. To understand the temporal gene expression of *catDE*, we monitored the Fur occupancy on this operator site using a chromosomal FLAG-tagged Fur in both WT and P_*spac*_-*frvA*. Using ChIP-qPCR, we found that Fur dissociated rapidly from the DNA binding site and ^~^50% decrease in Fur occupancy was observed within 3 min upon FrvA induction (Fig. 8). We then compared the Fur occupancy on this site with that on three Fur target sites that are representatives of the three sets of early, middle, and late genes determined previously (3). The results demonstrated that Fur occupancy on the *catDE* operator site followed the same pattern as that on the operator site of the late gene *fsrA* (Fig. 8), suggesting that the *catDE* expression is induced after derepression of BB biosynthesis and BB-mediated uptake systems. We infer that Fur derepression of *catDE* likely occurs soon after the onset of BB synthesis, and this will lead to an initial modest increase in *catDE* expression that will preemptively protect cells against the ensuing import of BB or other catecholate siderophores. In addition, because of the cooperative interaction of Fur and CatR *in vivo*, the loss of Fur repression will also render the CatR repressor more sensitive to accumulating catechols. Together, the loss of Fur and CatR repression will enable the effective detoxification of siderophore-derived catechol compounds.

**Fig. 8.**
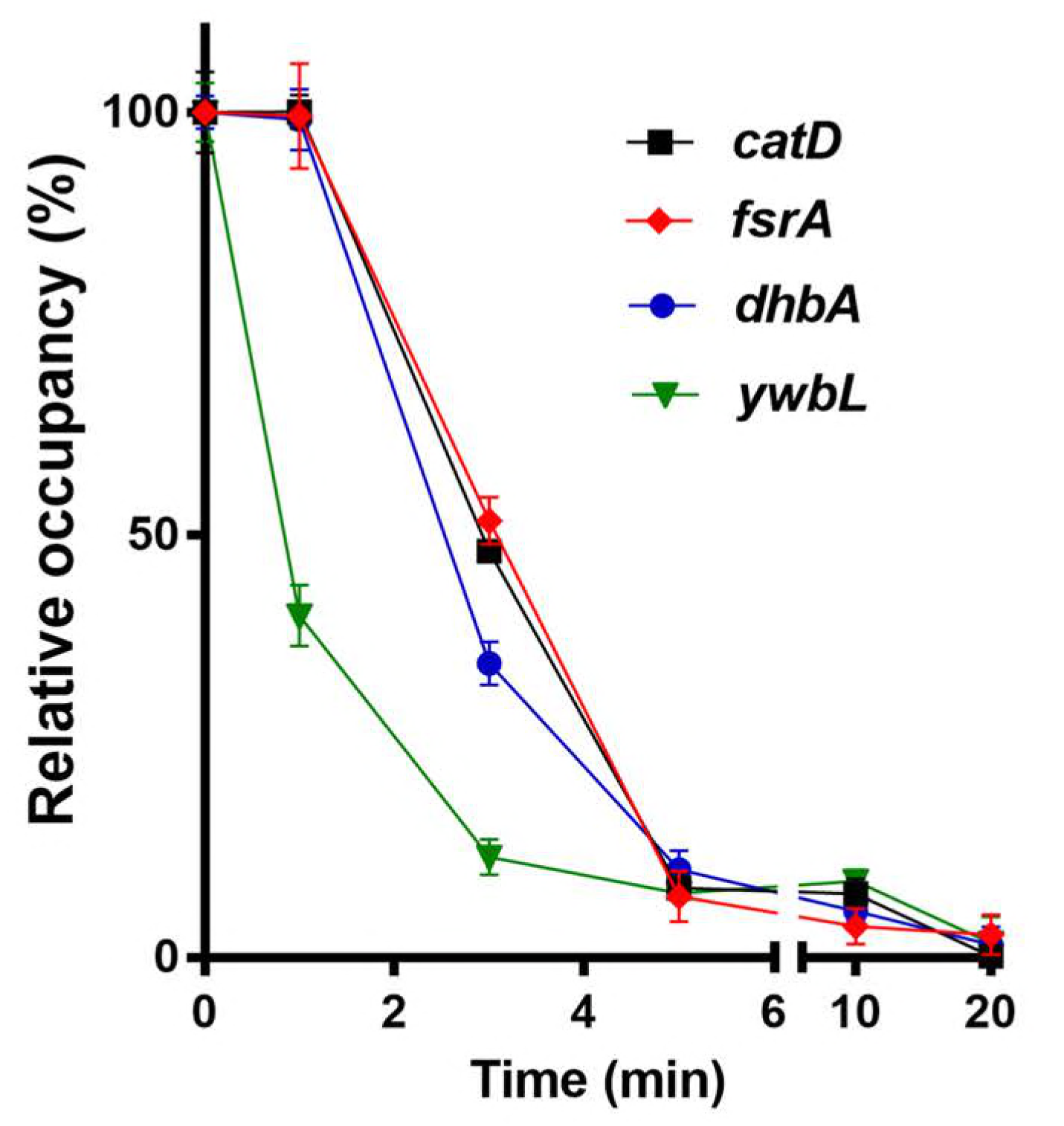
Fur occupancy on different operator sites. Fur occupancy was evaluated by ChIP-qPCR. DNA enrichment was calculated based on the input DNA (1% of total DNA used for each ChIP experiment). Fur occupancy at different sites was set as 100% at zero time-point. Dates are presented as the relative percentage (%) of occupancy at different time points after 1 mM IPTG induction of FrvA (mean ± SEM; n = 3). No significant DNA enrichment was observed for *gyrA* that is used as a nonspecific negative control.

### Concluding Remarks

Here we have provided a global overview of potential targets of Fur-mediated gene regulation by mapping of Fur binding sites under iron-replete conditions and after the onset of iron deprivation. Our work confirms the core Fur regulon as defined previously (9), and additionally suggests several new targets deserving of further study. These include potential roles for Fur in regulation of the *S477*-*ykoP* and *pps* (plipastin synthesis) operons, and in expression of FapR (a regulator of fatty acid synthesis), GidA (a tRNA modifying enzyme), CspB (cold-shock protein) and NarJ (nitrate reductase). Here, we focused our attention on the role of Fur in regulation of *catDE*, encoding a catechol 2,3-dioxygenase.

Catechol 2,3-dioxygenase is an exceptionally well-studied enzyme notable for its central role in the biodegradation of a wide variety of aromatic compounds that generate catechol intermediates. Here, we provide a novel example of an endogenously produced intoxicant that relies on CatDE for its degradation (Fig. 9). After import for ferric-bacillibactin into the cytosol, the BesA esterase cleaves the triscatecholate siderophore to release iron yielding three molecules of the BB monomer, 2,3-dihydroxybenzoate-Gly-Thr. In the absence of CatDE, this molecule, or its further degradation products, can be toxic and trigger cell lysis (Fig. 9). The expression of CatDE is under complex control involving three cooperatively functioning repressors. Binding of Fur to the *catDE* regulatory region appears to require cooperative interactions (largely with CatR). Upon the onset of iron deprivation there is an initial modest induction of *catDE* (as inferred from the effect of a *fur* mutation) that will preemptively synthesize CatDE. As catechol compounds accumulate in the cell, due to import and processing of ferric bacillibactin or import of other catecholate xenosiderophores, inactivation of the CatR repressor will lead to full induction. Since Fur and CatR bind cooperatively, once Fur is released the CatR repressor will be more easily inactivated, suggesting that the system will be poised for a rapid response. To our knowledge, this is the first example of an endogenous intoxicant that is catabolized by CatDE. We also present an example of a Fur target that is dependent on the cooperative action of multiple repressor proteins. This is reminiscent of the cooperative interactions documented previously for Fur, NsrR, and ResD reported for cells growing under anaerobic condition (15).

**Fig. 9.**
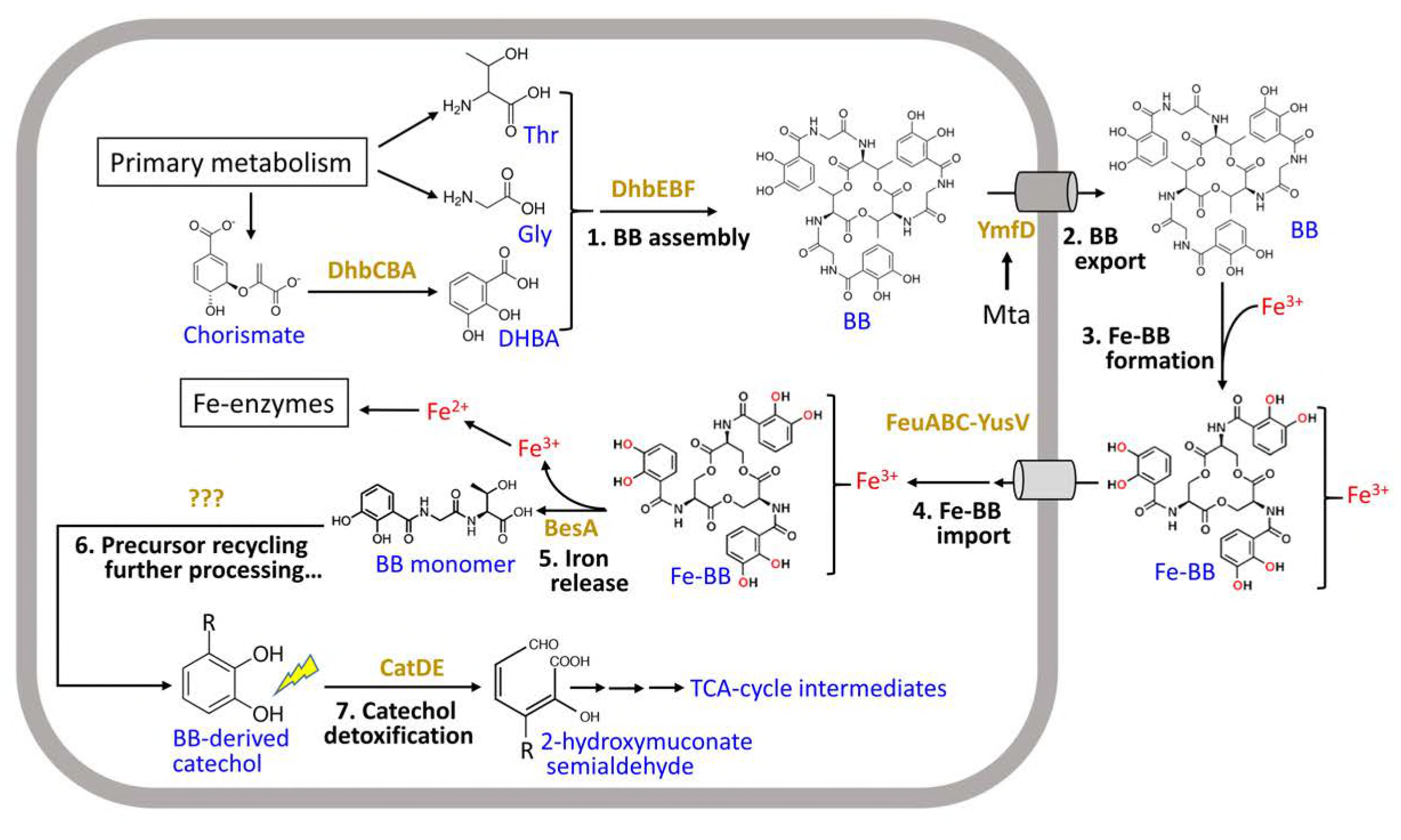
Correlation between BB metabolism and catechol detoxification. The endogenous siderophore BB is synthesized by an NRPS (non-ribosomal peptide synthetase) assembly system (DhbACEBF) (23) and secreted by a major facilitator superfamily transporter YmfD, which is under regulation of the transcriptional activator Mta, a MerR family regulator of multidrug-efflux transporter system (24). BB chelates iron with very high affinity and the resulting ferric-BB complex is then imported back into the cytosol through the FeuABC-YusV system and hydrolyzed by the BesA esterase to release iron (25), which yields three BB-monomers (2,3-dihydroxybenzoate-Gly-Thr). It is still unknown whether or how the BB monomer is further processed. Nonetheless, it is clear that the BB monomer and perhaps BB-derived catechol compounds require CatDE for detoxification during metabolism.

## Materials and Methods

### Bacterial strains and growth conditions

All strains used in the study are derivatives of *B. subtilis* CU1065 and are listed in <u>Table S1</u>. Cells were grown in LB or Belitsky minimal medium and growth was monitored using a Bioscreen growth analyzer as described in <u>Text S1</u>.

### RNA extraction and quantitative PCR (qPCR)

Cells were grown at 37°C in LB medium and RNA purified for qPCR analysis as indicated in <u>Text S1</u> using the indicated primers (<u>Table S2</u>).

### Disk diffusion assay

Cells were grown in Belitsky minimal medium and assayed for sensitivity to 10 μl of 1 M catechol or 200 mM dipyridyl as described in <u>Text S1</u>. The data are expressed as the diameter of the inhibition zone (mm).

### ChIP-seq, ChiP-qPCR and data analysis

*B. subtilis* cells expressing a C-terminal FLAG-tagged Fur at its native locus and an ectopic copy of *frvA* integrated at *amyE* locus were grown in LB medium amended with 25 μM iron to ensure Fur repression (2). 1 mM IPTG was added to induce expression of FrvA to deplete intracellular iron. ChIP was performed and analyzed by either Illumina-based sequencing (ChIP-seq) or qCR (ChIP-qPCR) as described in detail in <u>Text S1</u>.

### Electrophoretic mobility shift assays (EMSA)

Binding of Fur (activated with 1 mM MnC_2_) to the *catD* promoter region was monitored using an EMSA assay. The *K_d_* value, corresponding to the concentration of Fur that gives rise to 50% half-maximal shifting of the DNA probe, was evaluated and compared to *dhbA* as a positive control.

---

**Text S1. Materials and Methods**

**Table S1. Strains and plasmids used in this study**

**Table S2. Primer oligonucleotides**

**Table S3. Known Fur targets associated with ChIP-peaks.**

**Table S4. Putative Fur-regulated genes associated with ChIP-peaks**

**Table S5. Putative Fur target genes evaluated in this study**

**Fig. S1. putative Fur binding sites specifically under iron deplete conditions.**

Putative apo-Fur binding sites identified in the intragenic regions of: (A) *spo0B*, (B) *flhB*, (C) *xynP*, and (D) *ylpC* (also known as *fapR*). Two biological replicates were included as Exp.1 and Exp.2 for each condition. S/N denotes signal to noise ratio for peak calling.

**Fig. S2. CatDE is critical for catechol degradation.**

Sensitivity of the *sfp^0^* (A) and *sfp*^+^ strains (B) to catechol was evaluated in Belitsky minimal medium using a disk diffusion assay. 10 μl of 1 M catechol was applied to each disk. The data are expressed as the diameter (mean ± SEM; n = 3) of the inhibition zone (mm). Significant differences between wild type and mutant strains are indicated: ^∗∗^P < 0.01.

Representative growth curves in Belitsky minimal medium with *sfp^0^* (C) and *sfp*^+^ strains (D). 2mM catechol was used for both sets of experiments.

**Fig. S3. Catechol toxicity is diminished in LB medium**

Representative growth curves of WT and *catD* null mutant strains grown in LB medium without or with addition of 8mM catechol.

**Fig. S4. Metal-catechol complexes alleviates catechol intoxication**

Representative growth curves of *sfp*^+^ *catD* mutant cells grown in Belitsky minimal medium without or with addition of 2mM catechol. To evaluate the effects of metal-catechol complexes on catechol intoxication, different concentrations of metal salts were tested: (A) FeSO_4_, (B) CuSO_4_, (C) MnCl_2_, and (D) ZnCl_2_.

